# A speech envelope landmark for syllable encoding in human superior temporal gyrus

**DOI:** 10.1101/388280

**Authors:** Yulia Oganian, Edward F. Chang

## Abstract

Listeners use the slow amplitude modulations of speech, known as the envelope, to segment continuous speech into syllables. However, the underlying neural computations are heavily debated. We used high-density intracranial cortical recordings while participants listened to natural and synthesized control speech stimuli to determine how the envelope is represented in the human superior temporal gyrus (STG), a critical auditory brain area for speech processing. We found that the STG does not encode the instantaneous, moment-by-moment amplitude envelope of speech. Rather, a zone of the middle STG detects discrete acoustic onset edges, defined by local maxima in the rate-of-change of the envelope. Acoustic analysis demonstrated that acoustic onset edges reliably cue the information-rich transition between the consonant-onset and vowel-nucleus of syllables. Furthermore, the steepness of the acoustic edge cued whether a syllable was stressed. Synthesized amplitude-modulated tone stimuli showed that steeper edges elicited monotonically greater cortical responses, confirming the encoding of relative but not absolute amplitude. Overall, encoding of the timing and magnitude of acoustic onset edges in STG underlies our perception of the syllabic rhythm of speech.

---

Across human languages, speech comprehension relies on transforming continuous speech into discrete syllables. This poses a major computational challenge for the brain, as syllables are not separated by silent gaps or other known cues in natural speech. It is widely assumed that syllabic discretization relies on the amplitude envelope, which refers to the slow amplitude modulation of the speech signal (4-16 Hz)^1–10^. However, how cortical ensembles actually represent the speech envelope to extract syllables is largely unknown.

One prevailing model is that cortex contains an analog representation of the moment-by-moment fluctuations of the amplitude envelope. This interpretation stems from the well-documented neurophysiological correlation between cortical activity and the speech amplitude envelope ^11–20^. Alternatively, it has been suggested that cortex detects discrete acoustic landmarks. The most prominent candidate landmarks are the peaks in the envelope^9, 21,22^ or, as suggested from animal studies, rapid increases in amplitude (also called auditory onset edges)^23–26^. A fundamental question is whether the brain representation of the speech envelope is analog or discrete; and if discrete, which landmark is represented, and how linguistic information cued by this landmark allows syllabic discretization.

A challenge in understanding the neural encoding of the speech envelope is that amplitude changes are highly correlated with concurrent changes in phonetic content. One major reason is that vowels have more acoustic energy (sonority) than consonants^27^. Therefore, to know whether encoding is specific to amplitude modulations alone, or to the concurrent spectral content associated with phonetic transitions, it is necessary to use measure neural signals with a high spatial resolution.

Our goal was to determine the critical envelope features that are encoded in the non-primary auditory cortex in the human superior temporal gyrus (STG), which has been strongly implicated in phonological processing of speech. In particular, we asked whether STG neural populations encode instantaneous envelope values or detects a discrete landmark^19^. We then sought to determine how the encoded acoustic envelope features relate to the linguistically-defined syllabic structure of speech. Furthermore, we asked whether the encoding of the amplitude envelope is distinct from the processing of spectral cues found in consonants and vowels.

To address these questions, we used direct, high-density intracranial recordings from the cortical surface (electrocorticography, ECoG), whose high temporal and spatial resolution allowed us to distinguish between model alternatives. The high spatial resolution of ECoG allowed us to localize specific envelope encoding neural populations on STG and to distinguish them from neural populations encoding other temporal features, such as onsets, or acoustic-phonetic features^28^. Determining how syllables are neurally encoded may re-define the neurolinguistic understanding of how we perceive the syllabic rhythm of speech.

## Results

### Discrete events are extracted from the continuous speech envelope in bilateral STG

We asked whether neural populations in human STG represent the instantaneous moment-by-moment values of the amplitude envelope or whether they detect discrete acoustic landmarks in the speech envelope. We refer to an instantaneous representation as one that reflects the amplitude of the speech signal at each time point. We compared this to two independent models of encoding of prominent temporal landmarks: peaks in the speech envelope (peakEnv, Fig. 1A, black arrows) and peaks in the first derivative of the envelope (peakRate, Fig. 1A, purple arrows). Figure 1A shows the timing of each of these landmarks in a sample sentence, with peakRate preceding peakEnv landmarks within each cycle of the envelope (between two consecutive envelope troughs). Both landmarks appear within each envelope cycle (i.e. envelope between two consecutive troughs), such that envelope cycle onset, peakRate and peakEnv events are equally frequent in speech (Fig. 1B). Note also that all three events occur as frequently as single syllables, a prerequisite for one of these events to server as a marker of syllables.

**Figure 1.**
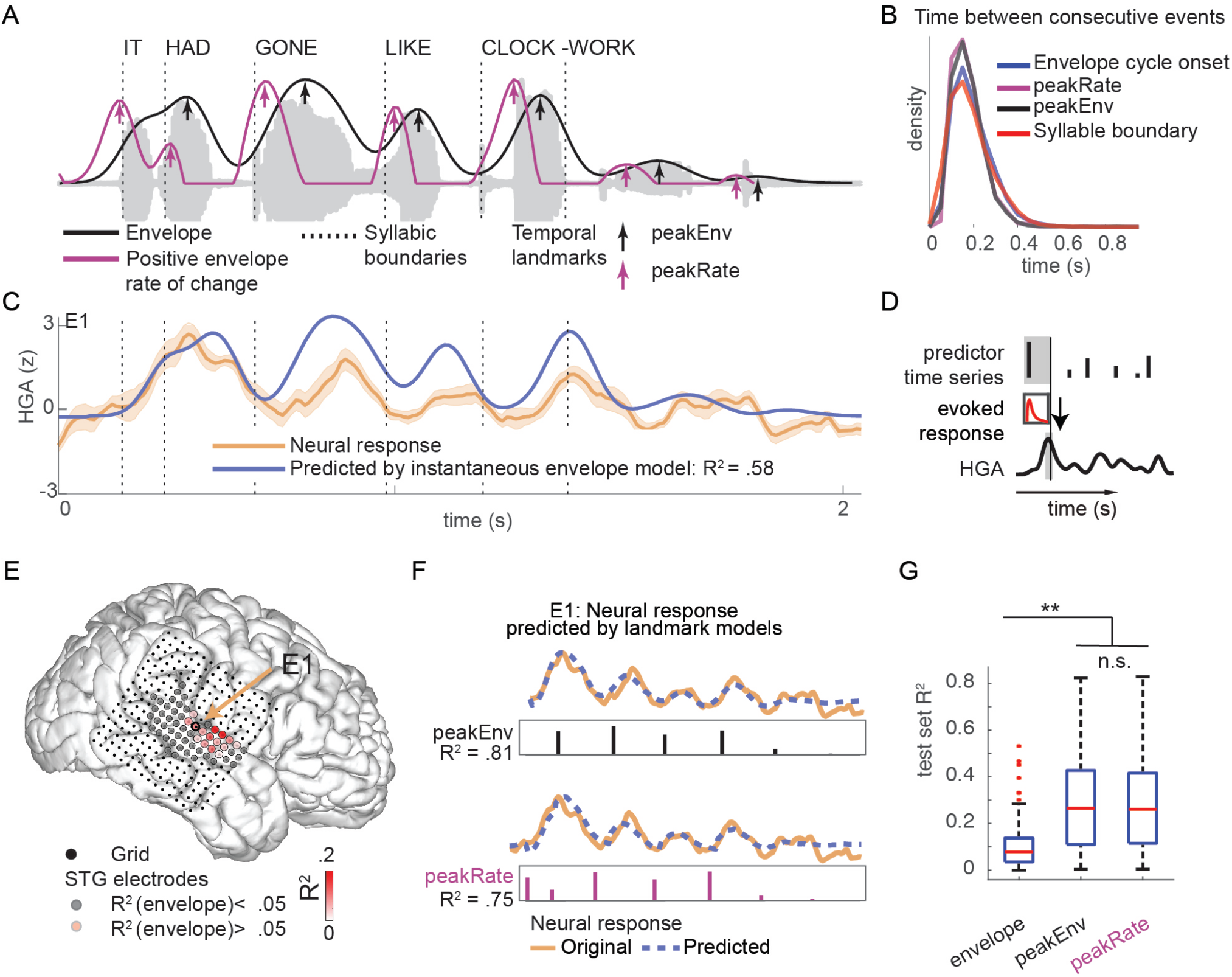
STG responses to speech amplitude envelope reflect encoding of discrete events. **A**. Acoustic waveform of example sentence, its amplitude envelope (black) and half-rectified rate of amplitude change (purple). Arrows mark local peaks in envelope (peakEnv) and rate of change of the envelope (peakRate), respectively. **B**. Rate of occurrence of syllabic boundaries, envelope cycles, peaks in the envelope, and peaks in the rate of change of the envelope in continuous speech across all sentences in stimulus set. All events occur on average every 200ms, corresponding to a rate of 5 Hz. **C**. Average HGA response to the sentence in A for electrode E1 (yellow). The predicted response based on a time-lagged representation of the envelope (blue) is highly correlated with the neural response for this electrode E1 and the example sentence (R^2^ = 0.58). **D**. Schematic of temporal receptive field model. The neural response is modeled as convolution of a linear filter and stimulus time series in a prior time window. **E**. Variance in neural response explained by representation of instantaneous amplitude envelope in an example participant’s STG electrodes. Neural activity in a cluster of electrodes in middle STG follows the speech envelope. **F**. Predicted neural response to the example sentence, based on discrete time series of peakEnv events (top) and peakRate events (bottom), in electrode E1. Both discrete event models outperform the continuous envelope model shown in C. G. Mean R^2^ for the instantaneous envelope, peakEnv, and peakRate models. Error bars represent the standard error of the mean (SEM) across electrodes. Both discrete event models are significantly better than the continuous envelope model, but they do not significantly differ from each other.

We used high-density ECoG recordings from the lateral temporal lobe of 11 participants (4 left-hemisphere, see Table S1 for patient details), who were undergoing clinical monitoring for intractable epilepsy and volunteered to participate in the research study. Participants passively listened to 499 sentences from the Texas Instruments and Massachusetts Institute of Technology (TIMIT) acoustic-phonetic corpus^29^, naturally produced by 402 male and female talkers (see Fig. 1A for example sentence). We extracted the analytic amplitude of neural responses in the high-gamma range (HGA, 70-150 Hz), which is closely related to local neuronal firing and can track neural activity at the fast rate of natural speech^30, 31^.

To compare the three models of envelope encoding, we first tested how well neural responses could be predicted from each model. For the instantaneous envelope model, we used the standard approach of cross-correlating neural activity and the speech envelope to determine the optimal lag at which the neural response resembles the speech envelope most closely. To model the neural data as a series of evoked responses to peakEnv or peakRate events, we used time-delayed multiple regression (also known as temporal receptive field estimation, Fig. 1D)^32, 33^. This model estimates time-dependent linear filters that describe the neural responses to single predictor events. All models were trained on 80% of the data and subsequently tested on the remaining 20% that were held-out from training. Model comparisons were based on held-out test set R^2^ values. For the comparison between models we excluded sentence onsets, because they induce strong transient responses in posterior STG after periods of silence typically found at the onset of a sentence or phrase, but do not account for variance related to the ongoing envelope throughout an utterance^28^.

In a representative electrode E1, HGA was well correlated with the speech amplitude envelope (across test set sentences: R^2^_mean_ = .19, R^2^_max_ = .66, mean lag = 60ms, Fig. 1C), but prediction accuracies were significantly higher for the landmark models (peakEnv model: R^2^_mean_ = .39, R^2^_max_ = .88; peakRate model: R^2^_mean_ = .39, R^2^_max_ = .84, Fig. 1F). This pattern held across all speech-responsive STG electrodes (see Figure 1E for electrode grid of a representative patient, Table 1 for model R^2^). Namely, HGA in up to 80% of electrodes was correlated with the speech envelope (n = 175 electrodes with sign-rank test p < .001, 3-34 per patient, average optimal lag: +86ms, SD = 70ms). However, landmark models outperformed the instantaneous envelope model (Table 1, signed rank test p’s < 10^−10^). Additional comparisons showed that both landmark models predicted the neural data equally well (signed rank test p > .5, Fig. 1G) and that a model with both sparse event predictors was not better than the models with only one landmark predictor (signed rank test p > .5). Moreover, other landmarks, such as troughs in the envelope and troughs in the rate-of-change of the envelope, predicted the data less well than the peakRate and peakEnv models (signed rank test p<.05) and did not explain additional variance beyond peakRate and peakEnv (signed rank test p>.5). In summary, these results demonstrate that the STG neural responses to the speech envelope primarily reflect discrete peakEnv or peakRate landmarks, not instantaneous envelope values.

**Table 1:** Model performance across all electrodes, in held-out stimulus set.

| *Model* | *Mean $R^2$* | *Max. $R^2$* |
| --- | --- | --- |
| *Continuous envelope* | .15 | .58 |
| *(cross-correlation)* |  |  |
| *peakEnv (evoked response)* | .30 | .82 |
| *peakRate (evoked response)* | .30 | .83 |

### Selective encoding of peakRate landmark revealed by slowed speech

Next, we wanted to understand which landmark – peakEnv or peakRate – is encoded in STG. However, at the natural speech rate, peakEnv and peakRate events occur on average within 60ms of each other (Fig. 2B). Thus, in natural speech, the encoding model approach used above could not disambiguate between them. To solve this, we created samples of slow speech that had longer envelope cycles (Fig. 2A and 2C) and thus also longer time windows between peakRate and peakEnv events (Fig. 2B). These sentences were still fully intelligible^34^ (see supplementary online materials for example sentences) and had the same spectral composition as the original speech samples (Fig. 2E). For example, in speech slowed to 1/4 of normal speed, the average time between consecutive peakRate and peakEnv events was 230ms, sufficient for a neural response evoked by peakRate to return to baseline before occurrence of the peakEnv landmark. Four participants listened to a set of 4 sentences that were slowed to 1/2, 1/3, and 1/4 of the original speech speed (Fig. 2A). We predicted that, in the context of the slowed sentences, evoked responses would be more clearly attributable to one of the envelope features (Fig. 2F).

**Figure 2.**
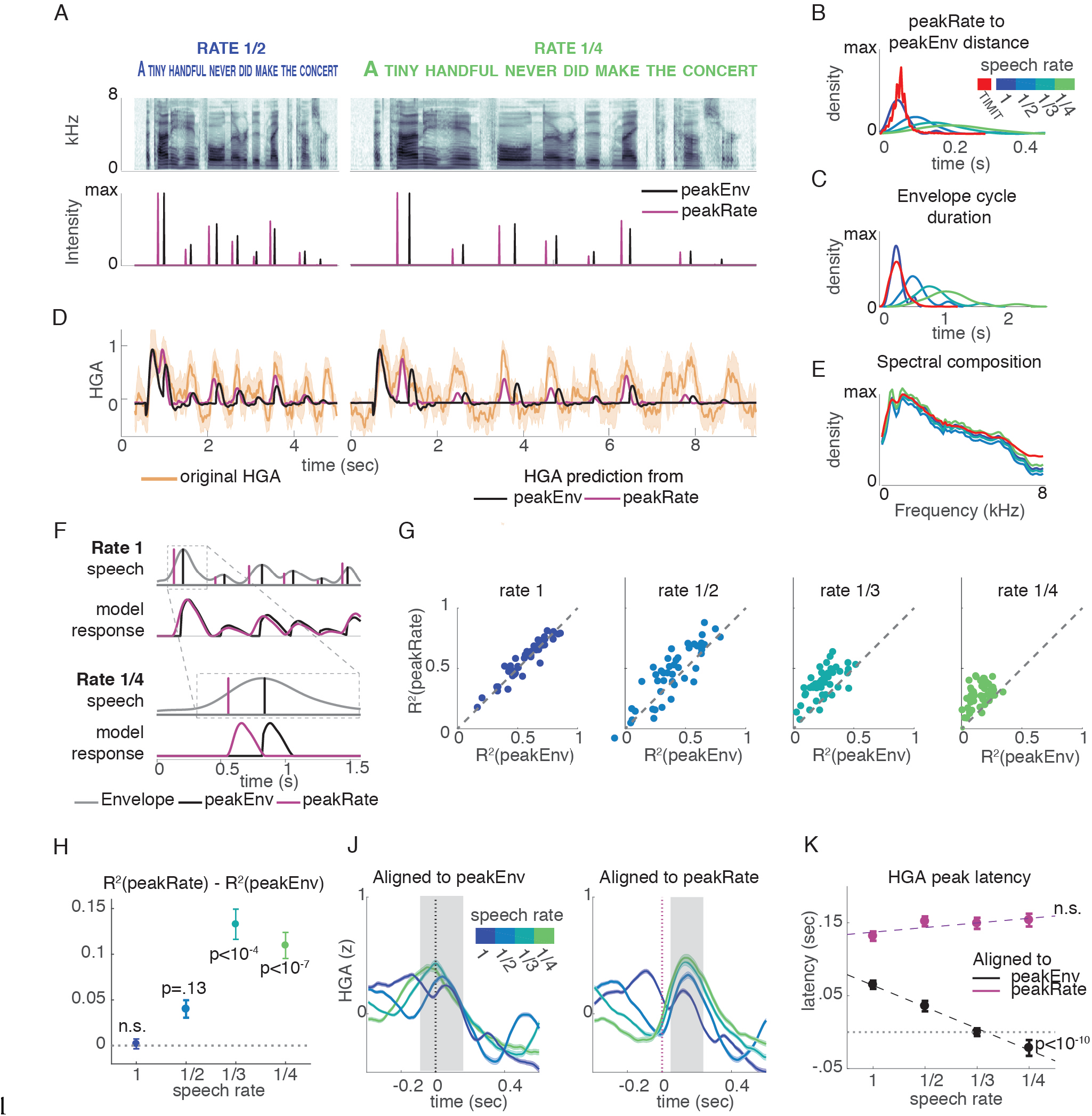
Neural responses to slowed speech demonstrate selective encoding of peakRate events. A. Top: Example sentence spectrogram at slowed speech rates 1/2 and 1/4. Bottom: Example sentence peakEnv and peakRate events for both speech rates. B. Distribution of latency between peakRate and subsequent peakEnv events, across all slowed speech task sentences and in full TIMIT stimulus set. Slowing increases time differences, and events become more temporally dissociated. C. Distribution of envelope cycle durations by speech rate, across all slowed speech task sentences and in full TIMIT stimulus set. Sentence slowing makes envelope cycles more variable, increasing discriminability. D. HGA response (orange) to an example sentence and neural responses predicted by sparse peakEnv (black) and peakRate (purple) models. Neural responses precede predicted responses of peakEnv model but are aligned with predicted responses of peakRate model accurately. E. Average spectral composition is similar for stimuli at different speech rates and the full set of TIMIT stimuli. F. Predicted neural responses for tracking of peakEnv events (black) and peakRate events (purple) for normally paced speech (top) and for slow speech (bottom). At rate 1, the models are indistinguishable. At rate 1/4, the models predict different timing of evoked responses. G. Comparison of test R2 values for peakRate and peakEnv models by speech rate in all speech-responsive STG electrodes. As speech rate is slowed, peakRate model explains neural responses better than peakEnv model. H. Mean (SEM) difference in R2 between peakEnv and peakRate models. The peakRate model significantly outperforms the peakEnv model at 1/3 and ¼ rates. J. Average HGA after alignment to peakEnv (left panel) and peakRate (right panel) events. Gray area marks window of response peaks across all speech rates, relative to event occurrence. When aligned to peakEnv events, response peak timing becomes earlier for slower speech. When aligned to peakRate events, response peak timing remains constant across speech rates. K. Mean (SEM) HGA peak latency by speech rate and alignment. Speech slowing leads to shortening of the response latency relative to peakEnv events only, such that it occurs before peakEnv events at the slowest speech rate.

Figure 2D shows neural responses to a single sentence at different speech rates for an example electrode, alongside with predicted responses based on the peakEnv and peakRate models. Neural responses at this electrode had the same number of peaks across speech rates, corresponding to single envelope cycles. At 1/2 rate, predictions from both models were almost identical, whereas at 1/4 rate a distinct lag between the predictions was readily apparent. Specifically, the predicted responses based on the peakEnv model lagged behind both the predictions from the peakRate model and the neural response.

Across all speech-responsive electrodes (n = 44, 8-25 per participant), we found that both models performed equally well at original and 1/2 speech rate, but that with additional slowing, the peakRate model became increasingly better than the peakEnv model (linear effect of speech rate: brate = .03, SE = .004, t (490) = 6.6, p<10^−10^, Fig. 2G and 2H).

In addition to comparing the model predictions, we also examined the average evoked responses aligned to peakEnv and peakRate events at different speech rates. We predicted that responses should be reliably time-locked to the preferred landmark event at all speech rates. The average responses across electrodes aligned to peakEnv events (Fig. 2J, left panel) reveal a neural peak that shifted backwards with speech slowing. Crucially, when speech was slowed by a factor of 3 or 4, neural HG peaks occurred concurrent with or even prior to peakEnv events (test against 0: rate3: b = 0, p = 1; rate 4: b = -.02, SE=.01, t (39) = 2.02, p = .05), providing clear evidence against encoding of the peakEnv landmark. In contrast, when aligned to peakRate, neural responses peaked at the same latency at all speech rates (Fig 2J, right panel), as summarized in Fig. 2K (interaction effect between speech rate and alignment: F (1,341) = 19.7, p<10^−4^, effect of speech rate on alignment to peakEnv: F (1,167) = 55.8, p<10^−10^, effect of speech rate on alignment to peakRate: F (1,174) = 1.6, p= .2). Two further analyses supported this result. First, a comparison between the peakRate and an envelope trough (minEnv) model showed that peakRate events predicted neural data better than minEnv events (Figure S2). Second, a comparison of neural response alignments to peakRate vs. peakEnv in natural speech supported the peakRate over the peakEnv model (Figure S1). Moreover, the change in latency between acoustic and neural peaks also refutes the continuous envelope model, because this model assumes a constant latency between the acoustic stimulus and corresponding points in the neural response.

Taken together, the slow speech data show that STG neural responses to the speech amplitude envelope encode discrete events in the rising slope of the envelope, namely the maximal rate of amplitude change, and refute the alternative models of instantaneous envelope representation or evoked responses to peakEnv events.

### peakRate cues the phonological structure of syllables

Syllable units are considered the phonological building blocks of words and have significant influence on the rhythm of a language, including its prosody and stress patterns. Here, we aimed to understand how peakRate events, as acoustically-defined temporal landmarks, relate to the phonological structure of syllables.

In Figure 3A, we show an example sentence annotated linguistically for the components of a syllable unit: the onset and the rhyme (comprised of nucleus and coda)^35^. The onset is the consonant or consonant cluster that precedes the syllabic vowel nucleus, and the rhyme includes the vowel nucleus and any consonant sounds (coda) that follow. Because speech amplitude is highest during vowels^27^, we hypothesized that peakRate events would mark the transition between the syllable onset and its rhyme. (Note that the term ‘syllable onset’ here is distinct from our use of acoustic onsets described previously, which refer to the beginnings of sentences following long silences.)

**Figure 3.**
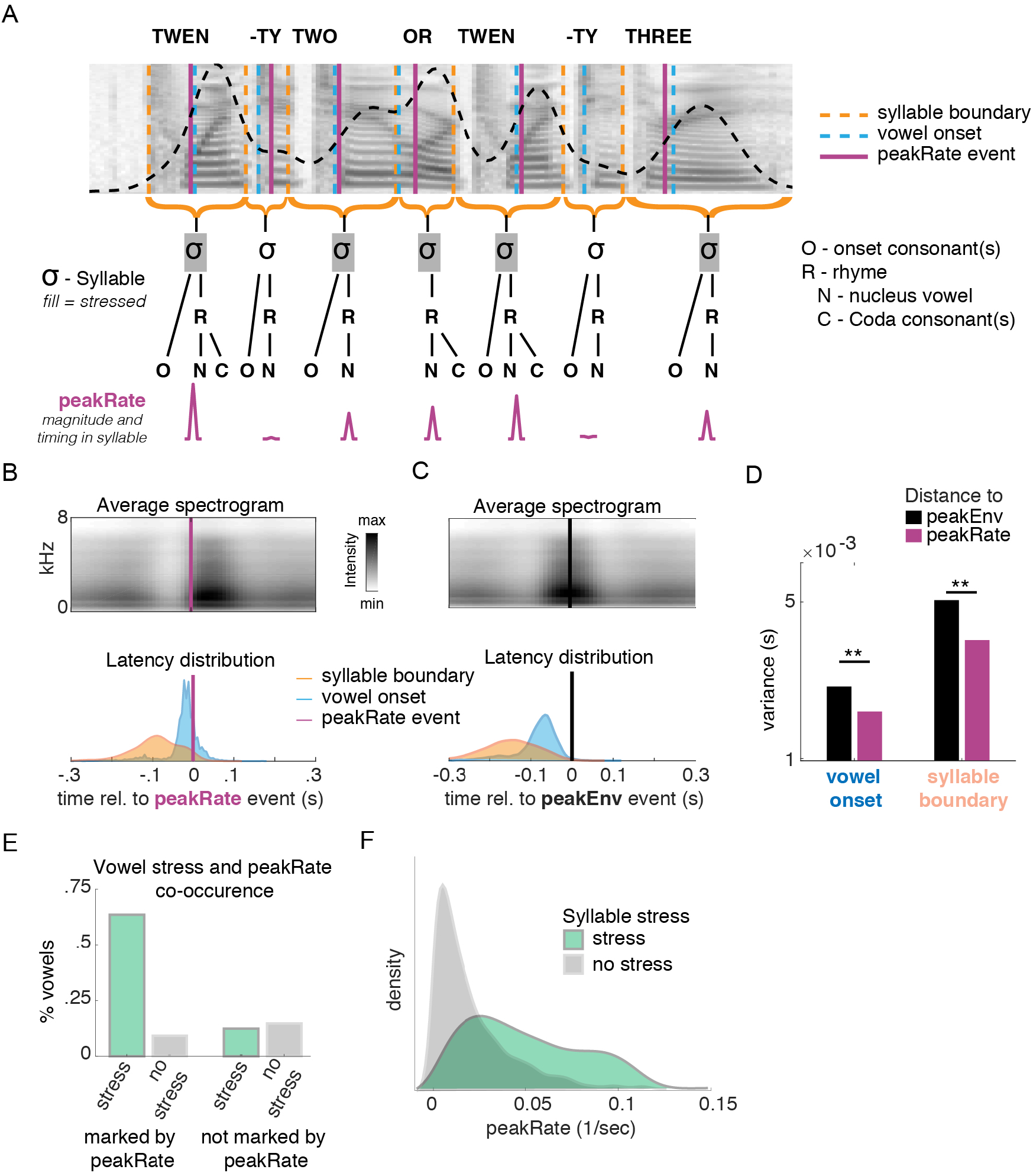
peakRate events cue the transition from syllabic onset consonants (C) to nucleus vowels (V). **A**. Upper panel: Spectrogram of an example sentence with syllabic boundaries, vowel onsets and peakRate events. peakRate events are concurrent with vowel onsets, but not with syllabic boundaries. Middle panel: Schematic of syllabic structure in the example sentence, marking stressed and unstressed syllables. Bottom panel: peakRate event location and magnitude within each syllable. PeakRate magnitude is larger for stressed syllables. **B, C**. Average speech spectrogram aligned to peakRate **(B)** and peakEnv **(C)** events. Top panel: Average speech spectrogram aligned to discrete event. peakRate events occur at time of maximal change in energy across frequency bands, whereas peakEnv events occur at times of maximal intensity across frequency bands. Bottom panel: Distribution of latencies of syllable boundaries and CV transitions relative to discrete event occurrence. CV transitions are aligned to peakRate events more than syllable boundaries. For peakEnv, both distributions are wider than for peakRate alignment. **D**. Variance in relative timing of syllable and vowel onsets and temporal landmarks. Smaller variance indicates that peakRate is a more reliable cue to vowel onsets that peakEnv. **E**. Co-occurrence of peakRate and vowels for stressed and unstressed syllables separately, in the TIMIT stimulus set. PeakRate is a sensitive cue for CV transitions, in particular to stressed syllables. **F**. Distribution of peakRate magnitudes in stressed and unstressed syllables. Above a peakRate value of .05 a syllable has a 90% chance of being stressed.

To test this, we analyzed the speech signal around peakRate events in the sentences in our stimulus set. In an example sentence in Figure 3A, the syllable /twen/ in the word ‘twenty’ has a delay between the syllable boundary and the peakRate event because of the consonant cluster /tw/, whereas the peakRate event is almost concurrent with the vowel onset. Across sentences, sound intensity increased rapidly at peakRate events (Fig. 3B top), which was due to peakRate events occurring nearly concurrently with the linguistically defined transition from syllable onset to syllable nucleus (mean latency between peakRate and vowel nucleus onset mean = 23ms, SD = 46ms, Fig. 3B bottom). On the contrary, the latency between peakRate events and syllable boundaries was significantly larger and more variable (mean = 86ms, SD = 63ms, t = −64, p<.001, Fig. 3B).

In comparison, peakEnv events mark the syllabic nuclei that are cued by peakRate events, as they occur after the peakRate events within a vowel (Fig. 3C). PeakEnv is, however, significantly less precise in cueing the C-V transition than peakRate (latency between peakEnv and C-V transition: mean = 86ms, SD = 53ms, comparison to peakRate bootstrap p’s <.05 for difference in means and variances, Fig. 3D). PeakEnv events thus inform the syllabic structure of a sentence by marking syllabic nuclei but were not informative with regard to the internal onset-rhyme structure of syllables.

In addition to serving as a reliable temporal landmark for syllables, we also found that the magnitude of peakRate events was important for distinguishing between unstressed and stressed syllables. In English, stress is cued by a combination of amplitude, pitch, and duration cues, and carries lexical information (i.e. distinguishing between different word meanings such as in ínsight vs. incíte). Despite the frequent reduction of syllables in continuous natural speech, peakRate events marked more than 70% of nucleus onsets overall and more than 80% of stressed syllable nuclei (Fig. 3C). The magnitude of peakRate was larger for stressed syllables than for unstressed syllables (Fig. 3D, sensitivity: d’=1.06). PeakRate events thus provide necessary information to extract the timing of syllabic units from continuous speech, the critical transition from onset to rhyme within a syllable, and the presence of syllabic stress. While many theories have posited the role of envelope for syllabic segmentation, i.e. detecting syllable boundaries, our results provide neurophysiological evidence for the alternative that peakRate is a landmark for the onset of the syllabic vowel nucleus, the importance of which has been previously hypothesized for the cognitive representation of the syllabic structure of speech^36, 37^.

### Topographic organization of temporal feature encoding on STG

Previous research described the encoding of sentence and phrase onset from silence in the posterior STG and encoding of phonetic features, in particular vowel formants in ongoing speech in the middle STG^28, 38^. To understand how peakRate encoding fits within this global organization of the STG, and to identify whether encoding of peakRate is distinct from encoding of the spectral content of vowels, we fit the neural data with an extended time-delayed regression model that included binary sentence onset predictors, consonant phonetic feature predictors (plosive, fricative, nasal, dorsal, coronal, labial), and vowel formants (F1, F2, F3, F4), in addition to peakRate (Fig 4A).

**Figure 4.**
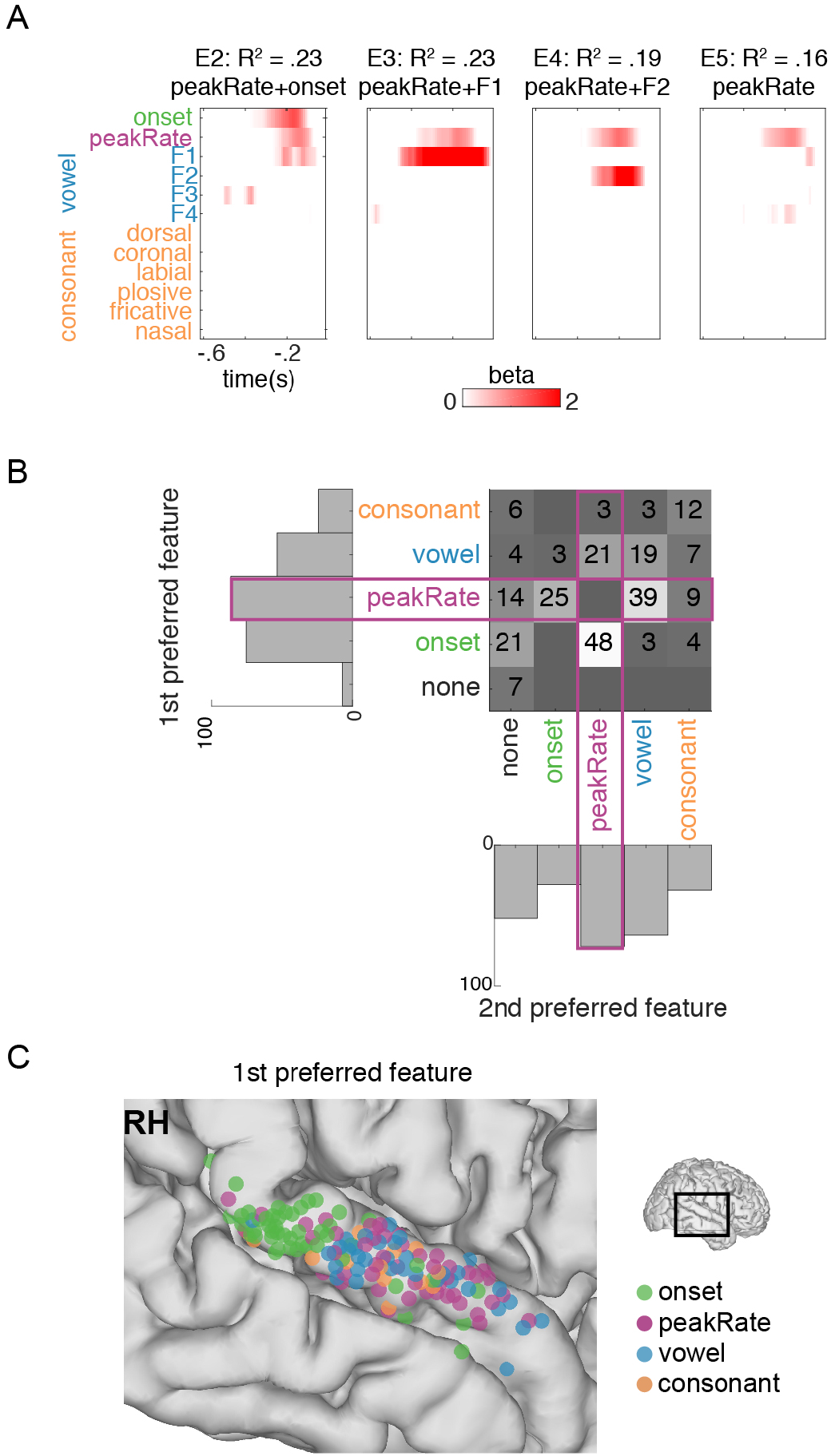
Independent and joint encoding of peakRate and other speech features. **A**. Linear weights from an encoding model with phonetic features and peakRate events for four example electrodes. Different electrodes show encoding of different features alongside peakRate. **B**. Number of electrodes with different combinations of the two significant features with the largest linear weights across STG electrodes. Vowel formant predictors (blue) and consonant predictors (orange) are each combined for visualization purposes. Onset and peakRate are blank along the diagonal because they contain one predictor only. peakRate encoding co-occurs with different phonetic features (e.g. E2-E4 in A) but can also occur in isolation (E5 in A). **C**. Anatomical distribution of electrodes with primary encoded onset, peakRate, vowel, or consonant features across all right hemisphere electrodes. Onset encoding is clustered in posterior STG, and peakRate encoding is predominant in middle STG.

We found that 80% of electrodes (E) significantly responded to at least two features and identified the two most preferred features for each electrode with various combinations of features present on single electrodes. Figure 4A shows feature receptive fields on four example electrodes with joint encoding of peakRate and onsets (E2), peakRate and first (E3) or second (E4) formant, and peakRate encoding only with marginal effects of onset and formant values (E5). Overall, peakRate was most frequently encoded by itself (n = 21 electrodes), or co-encoded with sentence onsets (n = 25+48 = 73 electrodes) or vowel formants (n = 21+39 = 60 electrodes, Fig. 4B).

Anatomically, encoding of peakRate was most prominent in middle STG (F (1,781) = 15.7, p < 10^−4^). This pattern was distinct from the anatomical distribution of onset responses, which were strongest in posterior STG (F (1,781) = 32.8, p < 10^−8^, Fig. 4C), consistent with our previous work^28^.

Taken together, these results indicate that distinct neural populations represent the temporal structure of speech, through encoding of peakRate and sound onsets, and its spectral content, through encoding of the spectral structure that corresponds to phonetic content^38^.

### Amplitude-rise dynamics alone drive neural responses to peakRate

Temporal and spectral changes in natural speech are inherently correlated. As a result, one potential confound is that peakRate encoding actually reflects the spectral changes that occur at the CV transition in syllables. We therefore asked whether STG responses reflect amplitude rise dynamics in the absence of concurrent spectral variation by using a set of non-speech amplitude-modulated harmonic tone stimuli (AM tones) in a subset of 8 participants.

AM tone stimuli contained amplitude ramps rising from silence (ramp-from-silence condition), or from a ‘pedestal’ at baseline amplitude of 12dB below the ramp peak amplitude (ramp-from-pedestal condition), as shown in Figure 5A. These two conditions were designed to broadly resemble amplitude rises at speech onset and within an ongoing utterance, respectively, but without any spectral modulations (such as vowel formant transitions) and variation in peak amplitude (such as amplitude differences between unstressed and stressed vowels) that are correlated with the amplitude rises in speech. Ramp durations and peak amplitude were kept constant across all stimuli, whereas rise times were parametrically varied (10-15 values between 10 and 740 ms, see table S1 for all rise time values) in both silence and pedestal conditions. The stimuli had complimentary rising and falling slopes, together ensuring equal stimulus durations. To simplify the analyses across the silence and pedestal conditions, we describe these stimuli in terms of amplitude rate-of-change ([peak amplitude – baseline amplitude]/rise time, i.e. amplitude rise slope, Fig. 5B). Because the amplitude rose linearly, the maximal rate of amplitude change (peakRate) occurred at ramp onset and was consistent throughout the rise.

**Figure 5.**
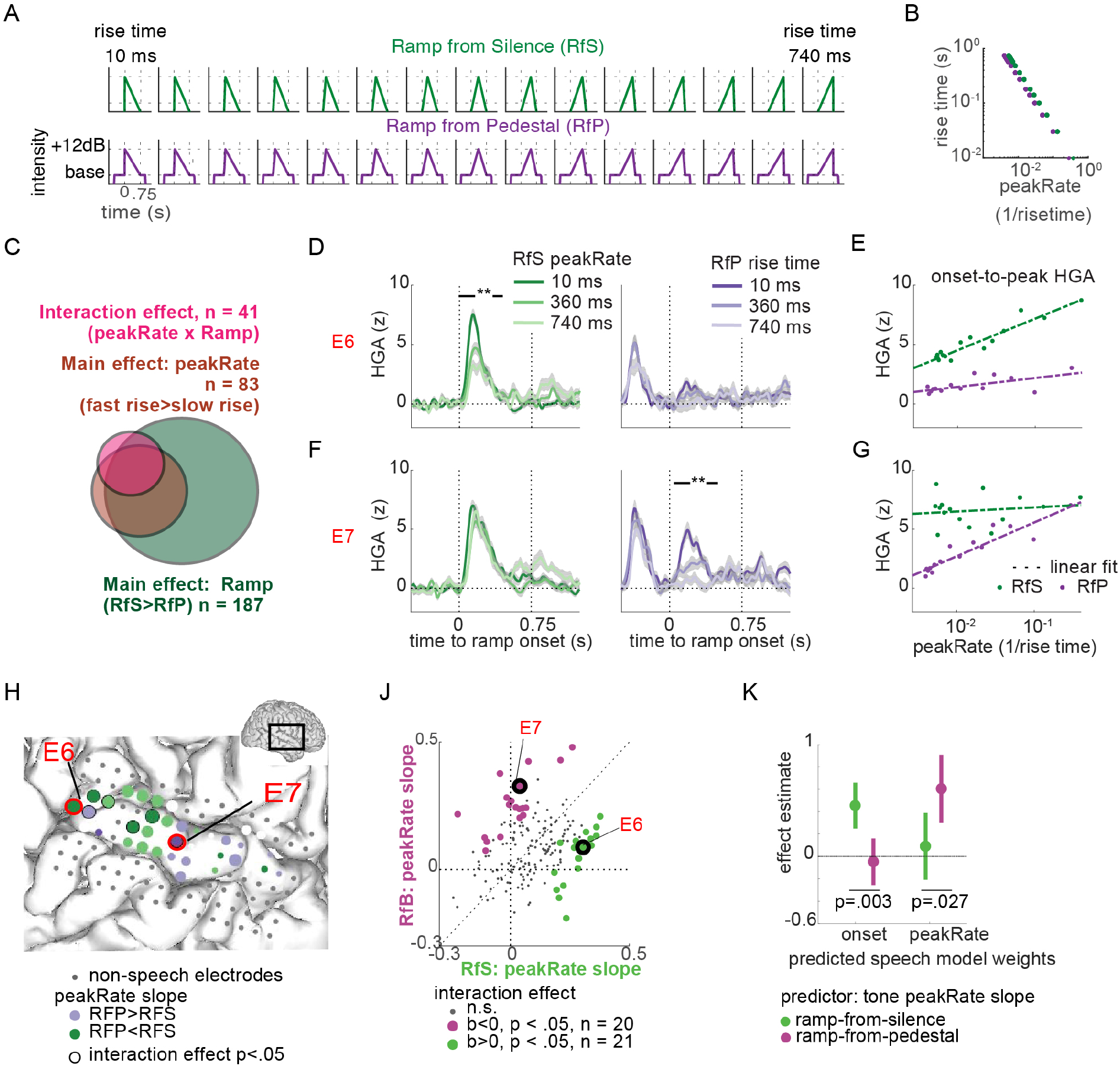
STG encoding of amplitude modulations in non-speech tones in onsets and in ongoing sounds. **A**. Tone stimuli used in the non-speech experiment. Tate of amplitude rise is manipulated parametrically, but peak amplitude and total tone duration is matched **B**. The relationship between ramp rise time and peakRate defined as for the speech stimuli. The peakRate value was reached immediately at ramp onset, as ramp amplitude rose linearly. **C**. Effect distribution across all electrodes. 20% of all electrodes showed a significant interaction effect between ramp type and peakRate, in addition to 78% showing a main effect of ramp type and 37% showing a main effect of peakRate. **D**. HGA responses to tones with three selected ramp rise times in ramp-from-silence (RfS; left) and ramp-from-pedestal (RfP; right) conditions in example electrode E6. **E**. Onset-to-peak HGA in electrode E6 as function of ramp peakRate, separately for RfS and RfP conditions. E6 codes for amplitude rate of change in RfS but not in RfP condition. **F**. Same as C. for example electrode E7. **G**. Same as D for example electrode E7. E7 codes for amplitude rate of change in RfP condition, but not in RfS condition. **H**. Temporal lobe grid from an example patient, with example electrodes E6 and E7 marked in red. Electrode color codes for relative magnitude of the peakRate effect on peak HGA in tone conditions. The purple electrodes’ HGA was more affected by peakRate in RfP condition, and the green electrodes’ HGA was correlated with peakRate values in RfS condition more than in RfP condition. Electrode size reflects maximal onset-to-peak HGA across all conditions. **J**. Slopes of peakRate effects on peak HGA, separately for each ramp condition. In colored electrodes the ramp condition x peakRate interaction was significant. Two distinct subsets of electrodes code for rate of amplitude change in one of the two conditions, only. **K**. Linear weights from a multiple regression model that predicted onset and peakRate linear weights in the speech model from peakRate slopes in tone model across electrodes. Representation of amplitude modulations at onsets and in ongoing sounds is shared in speech and in non-speech tones. Encoding of peakRate for envelope rises from silence is dissociated from peakRate encoding in ongoing sounds, in speech and in non-speech tone stimuli.

Analyses were focused on the same electrodes that were included in analyses of the speech task (n = 226 electrodes across 8 patients, with 11 – 41 electrodes per patient). Of these electrodes, 95% showed evoked responses to tone stimuli (false discovery rate (FDR)-corrected for multiple comparisons p < .05 for at least one of the effects in the ramp condition X rise time ANOVA analysis of peak amplitudes). Different rates of change were associated with differences in HG responses, which stereotypically started immediately after ramp onset and peaked at ~ 120ms. In particular, for stimuli with intermediate and slow rates of change, the neural HGA response peak preceded the peak amplitude in the stimulus (Figure S5A). This result further corroborates peakRate, and not peakEnv, as the acoustic event that drives neural responses^39^. Moreover, neural responses to ramp-from-pedestal tones returned to baseline between stimulus onset and ramp onset, despite an unchanged level of tone amplitude (sign rank test between HGA 0-200ms after stimulus onset and HGA 300-500ms after stimulus onset: p<10^−10^). This provides additional direct evidence for the encoding of amplitude rises and not the continuous envelope or amplitude peaks on STG.

### Distinct encoding of onsets and amplitude modulations in tone stimuli

Next, we wanted to test how the rate of amplitude rise would alter the magnitude of neural responses and whether neural responses would differentiate between preceding context, that is whether the ramp started from silence (analog to speech onsets) or from a pedestal (as in ongoing speech). We focused the following analyses on the effect of amplitude rise dynamics on the onset-to-peak magnitude of HGA responses, defined as the difference between the HGA at the time of ramp onset and at HG peak. We tested how peak HGA depended on the ramp condition (ramp-from-pedestal versus ramp-from-silence) and peakRate values by fitting a general linear model with predictors tone condition, peakRate, and their linear interaction, separately for each electrode.

Tone stimuli evoked robust responses in electrodes located in posterior and middle STG (see Fig. 5G for example electrode grid) with stronger responses to ramps starting from silence (mean beta = .3 across 187 out of 226 (82%) electrodes, which had a significant (p<.05) main effect of ramp condition, Fig. 5C). Moreover, on a subset of STG electrodes peak HGA was modulated by peakRate, with larger neural responses to fast rising ramps (mean beta = .2 on 83 out of 226 (37%) electrodes with significant (p<.05) main effect of peakRate, Fig. 5C). Similar to our findings in speech, some electrodes encoded peakRate in one of the two ramp conditions only, resulting in a significant interaction effect on 41 electrodes (18% of channels, χ^2^-test against chance level of observing the interaction effect on 5% of electrodes: z = 5, p < .01).

Electrodes E6 (Fig. 5D) and E7 (Fig. 5F) exemplify the two response patterns that drove this interaction effect, with a negative interaction effect in E6 and a positive interaction effect in E7. The amplitude of evoked responses in electrode E6 decreased with peakRate in the ramp-from-silence condition (b = .3, p < .05), but was not affected by peakRate in the ramp-from-pedestal condition (b = .08, p > .05; Fig. 5C right panel and Fig. 5E for peak HGA in all rise time conditions; linear interaction of ramp condition x peakRate: b = −.29, p < .05). Electrode E7 showed the opposite pattern, with a decrease in HGA for lower peakRate values in the ramp-from-pedestal condition (b = .32, p < .05, Fig. 5F right panel), but no effect of peakRate in the ramp-from-silence condition (b = 0.04, p > .05, Fig. 5F left panel, Fig. 5G for peak HGA in all rise time conditions; linear interaction of ramp condition x peakRate: b = .21, p < .05). Overall, neural activity on electrodes with a negative interaction effect (n = 20, green in Fig. 5J) encoded peakRate in the ramp-from-silence condition but not in the ramp-from-pedestal condition, whereas electrodes with a positive interaction effect (n = 21, purple in Fig. 5J) encoded peakRate in the ramp from pedestal condition only.

These results demonstrate that neural populations on STG encode amplitude rises independent from other co-occurring cues in speech. By parametrically varying peakRate in isolation from other amplitude parameters these data strongly support the notion that the STG representation of amplitude envelopes reflects encoding of discrete auditory edges, marked by time points of fast amplitude changes. These data also revealed a striking double-dissociation between the contextual encoding of peakRate in sounds that originate in silence and the encoding of peakRate in amplitude modulations of ongoing sounds, which indicates that dedicated neural populations track onsets after silences, e.g. sentence and phrase onsets, and intra-syllabic transitions.

### Amplitude rate-of-change encoding is similar in speech and non-speech tones

In a final analysis we tested whether encoding of peakRate events in non-speech tones reflects the same underlying computations as detection of peakRate events in speech. We reasoned that if neural populations encoded amplitude rises in tones and speech stimuli similarly, neural responses on electrodes that preferentially encode the dynamics of amplitude rises for amplitude ramps that start in silence (e.g., Fig. 5C, D) would also respond to sentence onsets in speech (as indicated by high beta values for the onset predictor in the speech encoding model). Conversely, we expected that electrodes that encode the dynamics of amplitude rises for ramps in ongoing tones (ramp-from-pedestal condition; e.g., Fig. 5E, F) would also encode peakRate events within sentences (as indicated by high betas for peakRate in the speech TRF model). To test this, we assessed whether speech model betas for onset and peakRate could be predicted from the same electrodes’ peakRate betas in the ramp-from-silence and ramp-from-pedestal conditions. Two separate linear multiple regressions were fit to predict speech model betas from the peakRate beta values in the tone task (Fig. 5K).

We found that responses to sentence onsets in speech were significantly predicted by encoding of peakRate in tone ramps starting from silence (b = 1.04, SD = .24, p = 10^−5^), but not by tracking of peakRate in tone ramps within ongoing tones (b =0.02, SD = 0.26, p = .9) and this difference was significant (permutation test of regression estimate equality, p = .003). Likewise, encoding of peakRate events after sentence onset in speech was not related to encoding of tone amplitude rise from silence (b = .04, SD = 0.17, p = .8), but it was significantly predicted by encoding of amplitude rise dynamics in ongoing tones (b =0.7, SD = 0.18, p = 10^−3^). Crucially, this difference was also significant (permutation test of regression estimate equality, p = .027). This analysis shows a robust overlap between the neural computations underlying the tracking of sound onset and amplitude modulation dynamics in speech and tones. Moreover, it corroborates the functional and anatomical dissociation between tracking of amplitude modulations in two distinct dynamic ranges – at onset and in ongoing sounds.

## Discussion

We show the amplitude envelope of speech is encoded by processing of peakRate landmarks in the middle STG. Neural responses scaled with peakRate magnitude, thus conveying not only the timing but also the velocity of envelope rises. Timing of this acoustically defined landmark marked the linguistic transition between syllabic onset and rhyme, and its magnitude indicated lexical stress. A further experiment with amplitude-modulated tones confirmed that neural responses to envelope rises encode the rate of amplitude change. Our decomposition of the speech envelope provides novel evidence for a discrete neural representation of envelope dynamics based on auditory edges, a flexible computational mechanism for encoding of the syllabic structure of speech across speech rates^5, 40^.

These findings have important implications for neuro-linguistic theories of speech processing. Previous work hypothesized that the amplitude envelope of speech enables the detection of syllabic units in continuous speech^6, 10,41^, but the actual computations that map the continuous envelope onto discrete syllabic units remained unclear. Our results provide a simple candidate mechanism to achieve two goals of syllable processing: 1) detection of timing and stress across syllables in an utterance, and 2) detection of the critical onset-rhyme transition within each syllable. Unlike previous suggestions^42^, peakRate events do not directly signal linguistically defined syllabic boundaries, which more closely correspond with troughs in the amplitude envelope^43^. However, listeners often have difficulty accurately perceiving syllabic boundaries, due to the fact that they are often not clear in natural speech because of large variation in how speech is produced, as well as theoretical disagreements about the placement of boundaries within consonant clusters^44, 45^. In contrast, there is strong behavioral evidence for the perceptual distinctiveness of an onset and rhyme in a syllable, and detection of a landmark at this transition may support this. For example, speech confusions often occur at the same syllable position (e.g. onsets are exchanged with other onsets)^37^

Several prominent theories of speech recognition have theorized that acoustic landmark events provide critical information for speech processing^46–48^. Amplitude rises are one such event, and indeed, the introduction of fast amplitude rises alone to vowel-like harmonic tones can induce the perception of a consonant-vowel sequence^49, 50^ and is critical for the correct perception of the temporal order of phonetic sequences^51, 52^. Our findings help to unify these seemingly disparate lines of speech research between acoustic and phonological theory.

The amplitude envelope is an important feature of sounds in general and amplitude envelope dynamics are encoded throughout the auditory system of different model organisms^53^. Single unit recordings along the auditory pathway up to secondary auditory cortices showed that the timing of single neural spikes and their firing rate reflect the dynamics of envelope rises^39,54–59^. Envelope encoding in human STG possibly emerges from amplitude envelope representations at lower stages of the auditory system. It thus may not be unique to speech processing, but rather is a universal acoustic feature with a direct link to the linguistic structure of speech and a crucial role in the comprehension of continuous speech.

Distinct neural populations in posterior STG encoded onsets (following silence), whereas those in the middle STG encoded acoustic edges in the ongoing utterance. Together, these populations encode envelope rises in distinct temporal contexts for sentences/phrases and syllables, respectively. Local neural populations within each zone can be tuned to specific phonetic features^38^. In summary, our results establish a cortical map for the temporal analysis of speech in the human STG.

## Methods

### Participants

11 (2 female) patients were implanted with 256-channel, 4mm electrode distance, subdural electrocorticography (ECoG) grids as part of their treatment for intractable epilepsy. Electrode grids were placed over the peri-Sylvian region of one of patients’ hemispheres (5 left, 6 right hemisphere grids). Grid placement was determined by clinical considerations. Electrode positions were extracted from post-implantation computer tomography (CT) scans, co-registered to the patients’ structural MRI and superimposed on 3-D reconstructions of the patients’ cortical surfaces using a custom-written imaging pipeline^60^. All participants were left-dominant and had normal hearing. The study was approved by the UC San Francisco Committee on Human research. All participants gave informed written consent prior to experimental testing. All patients participated in the speech experiment, a subset of 3 patients participated in the slow speech experiment, and a subset of 8 patients participated in the amplitude modulated tone experiment (Extended data table 1).

**Extended Data Table 1.** Patient details

| ID | Age | Gender | Handedness | Implant side | Seizure Foci | Resection Site | TIMIT natural speech | slow speech | AM tones |
| --- | --- | --- | --- | --- | --- | --- | --- | --- | --- |
| 1 | 33 | F | right | R | R Superior frontal gyrus | R Superior frontal gyrus | x |  | x |
| 2 | 28 | M | right | L | L Ant Temporal Lobe | L Ant Temporal Lobe | x |  | x |
| 3 | 37 | M | right | L | L Ant Temporal Lobe | L Ant Temporal Lobe | x |  | x |
| 4 | 20 | M | right | R | R Hippocampus | No resection (responsive neurostimulator) | x |  | x |
| 5 | 22 | M | right | R | R supramarginal gyrus | R Ant Temp Lobe | x |  | x |
| 6 | 40 | F | right | R | R Rolandic Cortex | No resection (responsive neurostimulator) |  |  | x |
| 7 | 36 | M | right | L | L Medial Temporal Lobe | No resection (responsive neurostimulator) | x |  | x |
| 8 | 59 | M | right | L | L Temporal Lobe | L ant Temporal Lobe | x |  | x |
| 9 | 24 | M | right | R | R Basal Temporal -Occipital lobe | R Post Temporal Lobe | x | x |  |
| 10 | 38 | M | right | L | L Temporal Lobe | L Amy/Hippocampus | x | x |  |
| 11 | 19 | M | right | R | R Frontotemporal Lobe | R Ant Temp and Frontal Lobe | x | x |  |

### Stimuli

All stimuli were presented at a comfortable ambient loudness (~ 70 dB) through free-field speakers (Logitech) placed approximately 80 cm in front of the patients’ head using custom-written MATLAB R2016b (Mathworks, https://www.mathworks.com) scripts.

### TIMIT speech

Participants passively listened to a selection of 499 English sentences from the TIMIT corpus (Ref), spoken by a variety of male and female speakers with different North American accents. Data in this task were recorded in 5 blocks of approximately 4 minutes duration each. Four blocks contained distinct sentences presented only once, and one block contained 10 repetitions of 10 sentences. This latter block was used for validation of temporal receptive field models (see below). Sentences were .9 – 2.4 s long and were presented with an inter-trial interval of 400ms.

### Slow speech

Slow speech stimulus set consisted of 4 sentences selected from the repetition block of the TIMIT stimulus set presented at 4 different speech rates: original, 1/2, 1/3 and 1/4. Participants listened to the stimuli in blocks of 5 minutes duration, which contained 3 repetitions of each stimulus with an inter-trial interval of 800ms. Each participant listened to 3-5 blocks of slow speech, resulting in 9-15 stimulus repetitions per participant.

### Amplitude modulated tones

In the non-speech tone task, participants passively listened to harmonic tones that contained an amplitude ramp starting either from silence (ramp-from-silence condition), or from clearly audible baseline amplitude (ramp-from-pedestal condition, Fig. 4A). The total duration of the amplitude ramp was 750ms. In the Ramp-from-pedestal condition, the ramp was preceded by 500ms and followed by 250ms of the tone at baseline amplitude (12dB below the peak amplitude). The peak amplitude of the ramp was the same across conditions. Ramp amplitude increased linearly from baseline/silence and then immediately fell back to baseline/silence for the remainder of the ramp duration. Ramp rise times could take 10-15 different values between 10 and 740ms (see Extended Data Table 2 for rise time values for every single patient). In the Ramp-from-silence condition the stimuli were harmonic tones with fundamental frequency 300 Hz and 5 of its harmonics (900, 1500, 2100, 2700, 3300 Hz). In the Ramp-from-pedestal condition half of the stimuli had the same spectral structure as the in the Ramp-from-silence background, and half the stimuli were pure tones of either 1500 or 2700 Hz. C-weighted amplitude was equalized between harmonics. Because neural responses to the ramp did not differ between harmonic and pure ramps we report all analyses pooled across these stimuli. Patients passively listened to 10 repetitions of each stimulus. Stimulus order was pseudo-randomized and the whole experiment was split into 5 equal blocks of approximately 5 minutes each.

For a comparison between conditions, we converted ramp rise times to rate of amplitude rise, calculated as

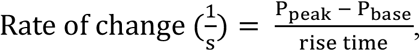

whereby P_peak_ and P_rise_ are sound pressure at ramp peak and at baseline, respectively. Because of the linear rise dynamics, the rate of amplitude rise reached its maximum at ramp onset and remained constant throughout the upslope of the ramp, so that peakRate was equal to the rate of amplitude rise.

**Extended Data Table 2.** AM-Tone task.

| Patient ID | Ramp from silence<br>Peak (dB rel. to ramp<br>pedestal amplitude) | Ramp from<br>background<br>Pedestal / amp. Diff. / Peak<br>(dB rel. to standard pedestal amplitude) | Tone in background |
| --- | --- | --- | --- |
| 1 | +12 | 0/12/+12 | 0/12/+12 |
| 2 | +12 | 0/12/+12 | 0/12/+12 |
| 3 | +12 | 0/12/+12 | 0/12/+12 |
| 4 | +12 | 0/12/+12 | 0/12/+12 |
| 5 | +12 | 0/12/+12 | 0/12/+12 |
| 6 | +12 | 0/12/+12<br>0/18/+18 | 0/12/+12 |
| 7 | +6 | 0/12/+12 | - |
|  | +12 | -6/12/+6 |  |
|  | +18 | -6/18/+12 |  |
| 8 | +6 | 0/12/+12 | - |
|  | +12 | -6/12/+6 |  |
|  | +18 | -6/18/+12 |  |

### Data analysis

All analyses were conducted in MATLAB R2016b (Mathworks, https://www.mathworks.com) using standard toolboxes and custom-written scripts.

### Acoustic feature extraction for speech stimuli

We extracted the amplitude envelope of speech stimuli using the specific loudness method introduced by Schotola^61^. This method extracts the envelope from critical bands in the spectro-temporal representation of the speech stimulus based on the Bark scale^62^ by square-rectifying the signal within each filter bank, band-pass filtering between 1 and 10 Hz, down-sampling to 100 Hz, and averaging across all frequency bands. We then calculated the derivative of the resulting loudness contours as a measure of the rate of change in the amplitude envelope. Finally, we extracted the sparse time series of local peaks in the amplitude envelope (peakEnv) and in its derivative (peakRate). This procedure resulted in a set of features for each cycle of the amplitude envelope (defined as the envelope between two neighboring local troughs, Fig. 1A inset): peakEnv and peakRate amplitudes, their latencies relative to preceding envelope trough, and the total duration of the cycle. Note that we did not apply any thresholding to definition of troughs or peaks, however we retained the magnitude of the envelope and its derivative at local peaks for all model fitting, such that models naturally weighted larger peaks more than small peaks. We also compared this envelope extraction method to a 10Hz low-pass filtered broadband envelope of the speech signal, which produced the same qualitative results throughout the paper.

### Neural data acquisition and preprocessing

We recorded electrocorticographic (ECoG) signals with a multichannel PZ2 amplifier, which was connected to an RZ2 digital signal acquisition system (Tucker-Davis Technologies, Alachua, FL, USA), with a sampling rate of 3052 Hz. The audio stimulus was split from the output of the presentation computer and recorded in the TDT circuit time-aligned with the ECoG signal. Additionally, the audio stimulus was recorded with a microphone and also input to the RZ2. Data were online referenced in the amplifier. No further re-referencing was applied to the data.

Offline preprocessing of the data included (in this order) down-sampling to 400 Hz, notch-filtering of line noise at 60, 120, 180 Hz, exclusion of bad channels, and exclusion of bad time intervals. Bad channels were defined by visual inspection as channels with excessive noise. Bad time points were defined as time points with noise activity, which typically stemmed from movement artifacts, interictal spiking, or non-physiological noise. From the remaining electrodes and time points, we extracted the analytic amplitude in the high gamma frequency range (70 – 150 Hz, HGA) using eight band-pass filters (Gaussian filters, logarithmically increasing center frequencies (70–150 Hz) with semi-logarithmically increasing bandwidths) with the Hilbert transform. The high-gamma power was calculated as the first principal component of the signal in each electrode across all 8 HG bands, using PCA. Finally, the HGA was down-sampled to 100 Hz and z-scored relative to the mean and standard deviation of the data within each experimental block. All further analyses were based on the resulting time series.

### Electrode selection

Analyses included electrodes located in the higher auditory and speech cortices on the superior temporal gyrus (STG), that showed robust evoked responses to speech stimuli, defined as electrodes for which a linear spectro-temporal encoding model^33^. explained a significant amount of variance (p<.001, see below for model fitting procedure, which was identical to the TRF fitting procedure). Analyses contained 227 electrodes, 3 – 34 within single patients.

### Analysis of the neural data in the speech task

#### General model fitting and comparison approach

All models were fit on 80% of the data and evaluated on the held-out 20% of the data set, as Pearson’s correlations of predicted and actual brain responses. There correlations were then squared to obtain R^2^, a measure of the portion of variance in the signal explained by the model. Model comparisons were conducted on these cross-validated R^2^ values, which were calculated separately for average neural responses (across 10 repetitions) for each test set sentence. Formal comparisons between R^2^ values across electrodes were conducted using Wilcoxon rank-sum test and a significance threshold of .05.

#### Representation of instantaneous amplitude envelope

To test whether neural data contain a representation of instantaneous amplitude envelope values, we calculated the maximum of the cross correlation between the speech amplitude envelopes and HGA, restricted to positive lags (i.e. neural data lagging behind the speech envelope). The optimal lag was determined on the training set of the data, model fit was then assessed on the independent training set (see above).

#### Time-delayed multiple regression model (TRF)

To identify which features of the acoustic stimulus electrodes responded to, we fit the neural data with linear temporal receptive field models (TRFs) with different sets of speech features as predictors. For these models, the neural response at each time point (HGA(t)) was modeled as a weighted linear combination of features (f) of the acoustic stimulus (X) in a window of 600ms prior to that time point, resulting in a set of model coefficients, b_1…, d_ (Fig. 1C) for each feature f, with d = 60 for a sampling frequency of 100 Hz and inclusion of features from a 600ms window.

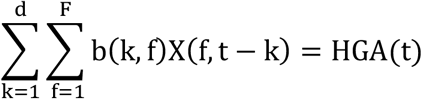

The models were estimated separately for each electrode, using linear ridge regression on a training set of 80% of the speech data. The regularization parameter alpha was estimated using a 10-way bootstrap procedure on the training data set for each electrode separately and then a final alpha value was chosen as the average of optimal values across all electrodes for each patient.

For all models, predictors and dependent variables were z-scored prior to entering the model. This approach ensured that all estimated beta-values were scale-free and could be directly compared across predictors, with beta magnitude being an index for the contribution of a predictor to model performance.

#### Feature receptive field models

To assess the extent of overlap between amplitude envelope tracking and phonetic feature encoding in STG electrodes, we also fit a time-delayed multiple regression model that included median values of the first four formants for all vowels, and place and manner of consonant articulation, in addition to onset and peakRate predictors. Phonetic feature and formant predictors were timed to onsets of the respective phonemes in the speech signal. Comparisons of beta-values for the different predictors were based on maximal beta values across time points. Phonetic features for this model were extracted from time-aligned phonetic transcriptions of the TIMIT corpus and standard phonetic descriptions of American English phonemes. Vowel formants were extrapolated using the freely available software package Praat^63^.

To assess the significance of predictors in TRF encoding models, we used a bootstrapping procedure. The model was refit 1000 times on a randomly chosen subset of the data. This was used to estimate distributions of model parameters. The significance of a single feature in the model was determined as at least 10 consecutive significant beta values for this feature (p < .05, with Bonferroni-correction for multiple comparisons across electrodes).

#### Spatial distribution test

The spatial organization of electrodes encoding onsets, peakRate, and phonetic features on STG was tested by predicting the beta values (b) for each feature from the location of electrodes along the anterior-to-posterior axis (p) of the MNI projection of single patients’ electrode location. To test for concentration of a feature in mid-STG, we included a quadratic term in the regression model:

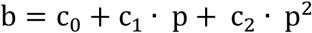

In this model, posterior-anterior effects of location onto beta values correspond to linear effects of p (i.e. c1 significantly different from 0). A concentration of significant beta values in mid-STG would correspond to c2 values above 0.

### Analysis of slow speech task

#### Temporal receptive field models

We tested model fits of the time-delayed multiple regression models that were fitted on the TIMIT training data for each of the four speech rate conditions. Note, that all 4 sentences that were presented in this task were part of the TIMIT test set. We used quality of model fits and the comparison between models at each speech rate as an indicator of whether STG retained a representation of the instantaneous amplitude envelope or of landmark events. We used a linear regression across electrodes to test whether the difference between peakEnv and peakRate model changed with speech rate.

#### Realignment to acoustic landmarks

Neural HGA data was segmented around peakEnv and peakRate landmark occurrence (400ms before and 600ms after each landmark) and averaged within each rate condition. The analysis included all landmark occurrences (n =21), excluding sentence onsets. We extracted the latency of HG peaks relative to both landmarks for each electrode. The effect of speech rate and landmark onto HGA peak latencies was assessed used a 2-way repeated-measures ANOVA with factor speech rate and landmark.

### Analysis of amplitude-modulated tone task

#### Data acquisition and preprocessing

Data acquisition and preprocessing followed the same procedure as for the speech data. However, z-scoring for the tone task was done separately for each trial based on HG mean and variance during the 500ms before stimulus onset. Responses were averaged across the repetitions of the same ramp condition and rise time combination before further analyses.

#### Responsiveness to tone stimuli and electrode selection

Because we were interested in characterizing how speech electrodes respond to non-speech amplitude modulated tones, analyses were performed on all electrodes that were included in the speech task. We quantified the response to ramp onsets as the trough-to-peak amplitude difference between the HGA in a 50ms window around ramp onset and the maximal HGA in the 750ms window after ramp onset.

#### General linear model of response amplitudes

We analyzed the effects of ramp type (ramp-from-silence vs. ramp-from-background) onto response trough-to-peak amplitude for every electrode separately, using a general linear model with predictors ramp type, log ramp rise time, and their linear interaction, with significance threshold set to p<.05 (uncorrected).

### Code availability

All custom-written analysis and stimulus presentation code is available upon request to the corresponding author (EC).

## Supplementary Figures

**Figure S1.**
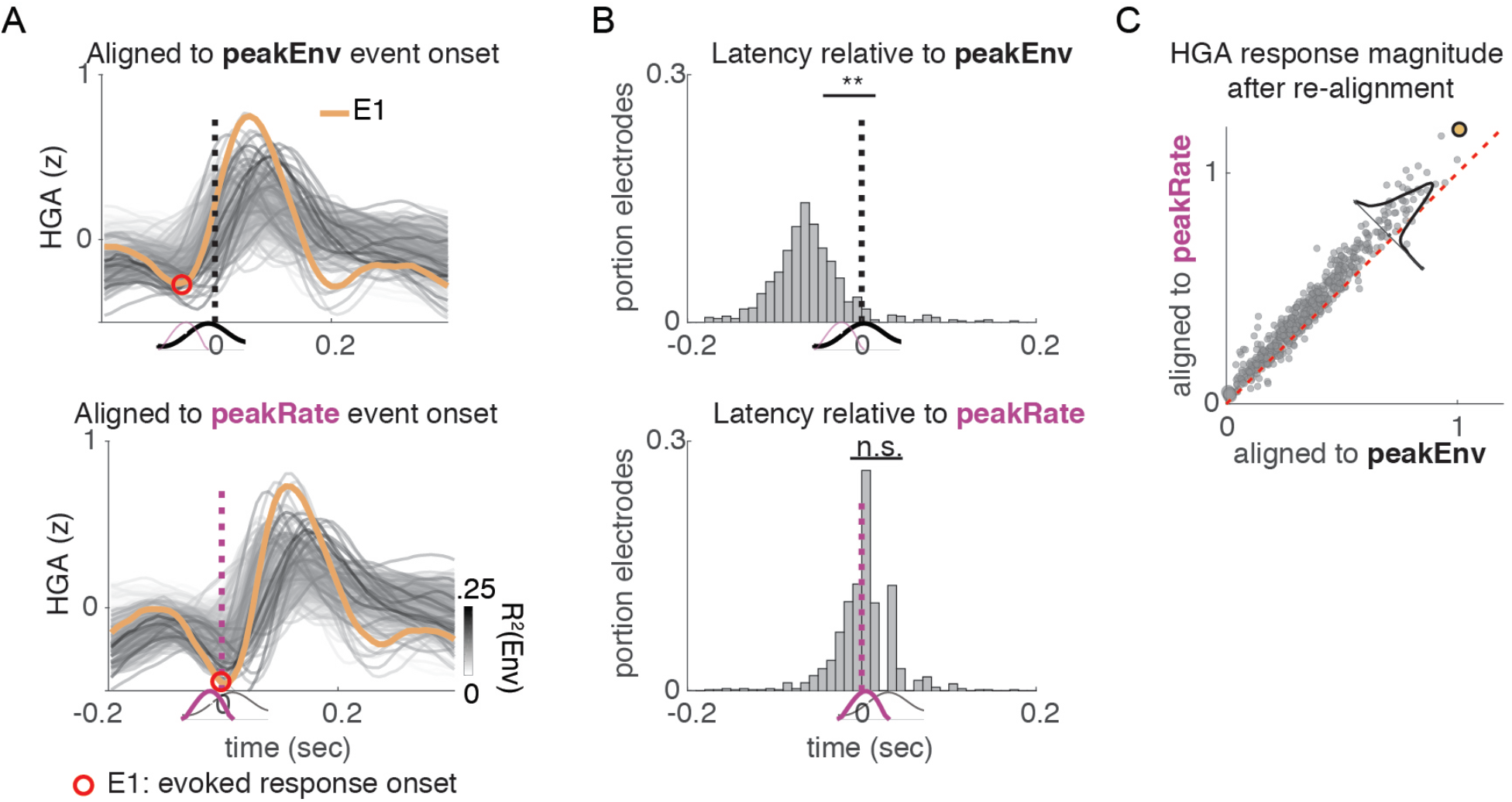
Segmentation of neural responses to naturally produced sentences (TIMIT) around peakEnv and peakRate events. **A**. Single electrode neural responses realigned and averaged across all peakEnv (top) or peakRate (bottom) events. Realignment shows that neural response onset is prior to peakEnv events but temporally aligned to peakRate events (bottom). **B**. Distribution of latency between trough in neural response and peakEnv (top) and peakRate (bottom) events. **C**. Magnitude of average neural response in alignment to peakEnv vs. peakRate. Response magnitude is larger for alignment to peakRate than to peakEnv. Latency and response magnitude analyses support peakRate encoding over peakEnv encoding.

**Figure S2.**
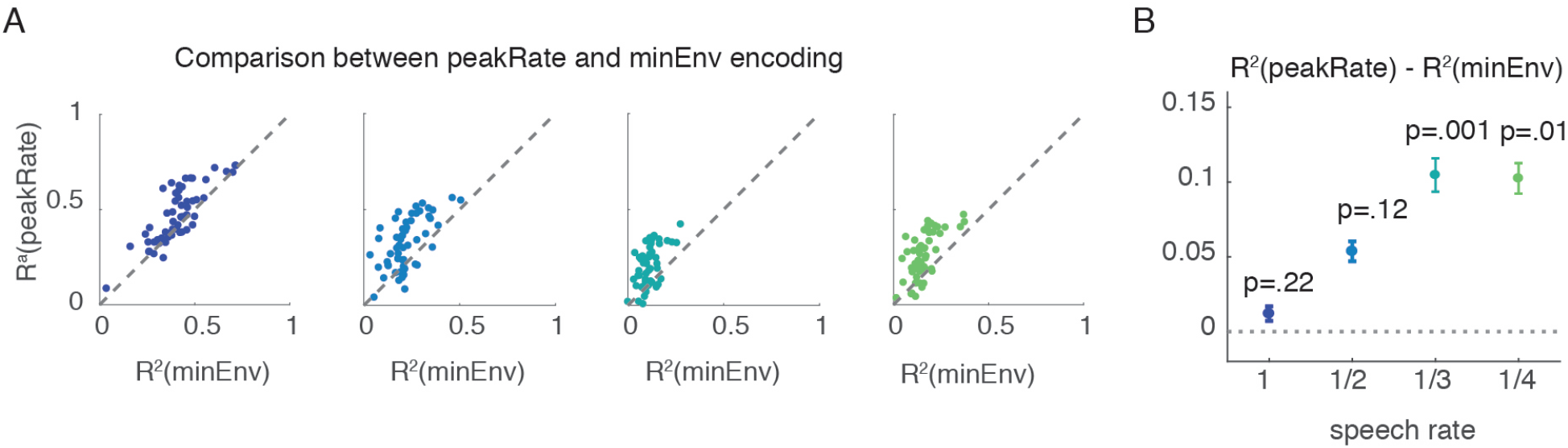
Comparison between neural response prediction based on peakRate and minEnv models. **A**. Across all electrodes, peakRate model outperformed minEnv model. **B**. Average difference in model R^2^ increases with speech slowing. PeakRate is thus a better predictor of neural activity than minEnv.

**Figure S3.**
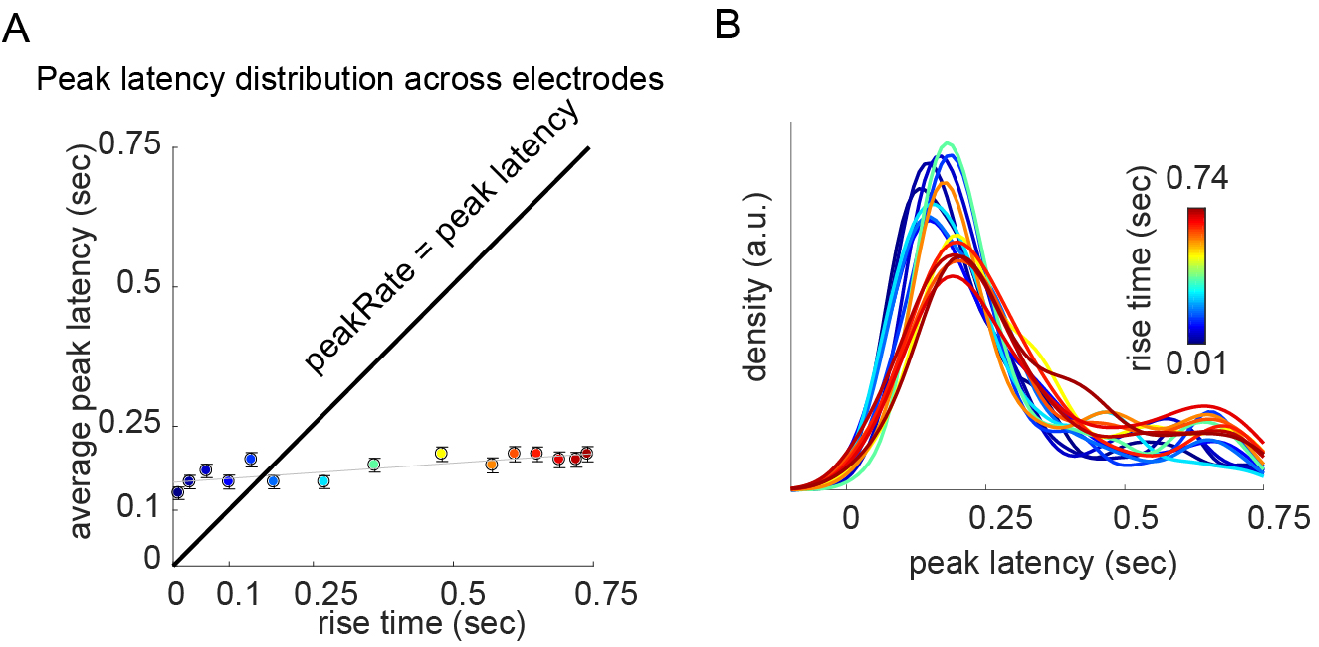
Latency of neural response peaks as function of ramp rise time in AM-tones. **A**. Peak latency as function of ramp rise time. For intermediate and long rise time the neural response peaks earlier than the ramp reaches maximal amplitude. **B**. Peak latency distribution across electrodes as a function of rise time.

## Acknowledgements

We thank K. Johnson, C. Schreiner, S. Nagarajan, S. Greenberg, A. Breska, L. Hamilton, J. Downer, P. Hullett, M. Leonard, B. Malone, and C. Tang for helpful comments and feedback. We also thank L. Hamilton and E. Edwards for providing some of their code for analysis of the TIMIT data and other members of the Chang Lab for help with data collection. This work was supported by grants from the NIH (R01-DC012379 to E.F.C and the German Research council (OG 105/1 to YO). E.F.C. is a New York Stem Cell Foundation Robertson Investigator. This research was also supported by the New York Stem Cell Foundation, the Howard Hughes Medical Institute, the McKnight Foundation, The Shurl and Kay Curci Foundation, and The William K. Bowes Foundation.

## Author contributions

Y.O. and E.F.C. conceived and designed the experiment. YO. and E.F.C. collected the data. Y.O. analyzed the data. Y.O. and E.F.C. wrote the paper.

## Corresponding author

Edward F. Chang

